# Spaniel: analysis and interactive sharing of Spatial Transcriptomics data

**DOI:** 10.1101/619197

**Authors:** Rachel Queen, Kathleen Cheung, Steven Lisgo, Jonathan Coxhead, Simon Cockell

## Abstract

Spatial Transcriptomics allows the sequencing of the complete transcriptomes from barcoded regions of intact tissue. The technology has the potential to answer a wide range of biological questions concerning cellular function, but analysis of the data presents a number of challenges which are not met by existing analysis tools. Here we present Spaniel, an R package providing a framework for analysing and sharing Spatial Transcriptomics data.

## Main

Whilst technologies such as single cell RNAseq provide tools for dissecting cellular heterogeneity, dissociation of the tissue mean that information about spatial relationship of the cells is lost. Spatial Transcriptomics (ST) ^1^ is a method for sequencing spatially barcoded transcriptomes from intact tissue sections. This provides a method of linking gene expression data to spatial location of a cell within a tissue cross-section, to further elucidate cellular function. Computational analysis of Spatial Transcriptomics data has a unique set of requirements which are not satisfied by existing analysis tools. Firstly, there is a strong visual element to the analysis and gene expression must be linked back to a histological image of the tissue. Secondly, to answer complex questions there is need for data integration both between different Spatial Transcriptomics datasets and with other datasets such as single cell RNAseq. Finally, a close collaboration between a computational biologist analysing the data and the wet-lab biologist with specialised knowledge of the tissue type is required to interpret the experiments, but as each Spatial Transcriptomics dataset generates a large amount of data and a number of decisions must be taken during analysis, it difficult to share data in a static manner.

Numerous tools exist for analysing single cell RNAseq and low input RNAseq experiments ^2^, but these lack methods for visualising histological data. The ST Viewer tool ^3^ has been designed for analysing and visualising ST data but as it is a standalone tool it is not suitable for integrative analysis of multiple datasets.

Here we introduce Spaniel, an R package which provides a framework for analysing and sharing Spatial Transcriptomics data. Spaniel provides methods for thorough quality control, visualisation and pre-processing of the data. It uses two pre-existing S4 objects designed for single cell analysis, namely Seurat object ^4^ and SingleCellExperiment object ^5^, providing strong potential for integration of and with other single cell datasets. Spaniel also provides a Shiny app which allows results to be shared in an interactive manner.

To test the Spaniel application and produce the figure presented here, we used publicly available sequencing data ^1^ from the mouse olfactory bulb and the hematoxylin and eosin (H&E), HE_Rep1 taken from the Spatial Transcriptomics website (http://www.spatialtranscriptomicsresearch.org/). Each Spatial Transcriptomics experiment involves sectioning a 5-16 μm slice of tissue which is placed on a slide containing a grid of spatially barcoded polyT probes. The probes are positioned in spots on the slide in a 35 * 35 grid, where each spot covers 10 – 100 cells. The sequencing data was generated using paired end sequencing where read 1 contains the spatial barcode and UMI and read 2 contains the transcript sequence. Spaniel takes a spatial transcriptomic expression matrix where each row corresponds to a gene and each column corresponds to a spot coordinate. To create the expression data, from the sequencing data, the paired FASTQ files were demultiplexed with a publically available perl script (https://github.com/tallulandrews/scRNASeqPipeline/blob/master/0_custom_undo_demultiplexing.pl) using the spatial barcodes encoded in read 1. Read 2 from successfully demultiplexed pairs were trimmed for quality using Trimmomatic version 0.36 ^6^. A reference was created using Ensembl mouse reference genome Release M20 (GRCm38.p6). The trimmed reads were aligned to this reference using STAR version 2.5.3a ^7^ in single read alignment mode. The number of reads were quantified using HTSEQ version 0.6.1 ^8^ and a count matrix was created. The H & E images were cropped to region around the edges of the spots and resized to 1000 × 1071 pixels with a resolution of 72dpi using a photo editor.

Spaniel includes a series of tools to aid the quality control and analysis of Spatial Transcriptomics data and is designed to be used alongside existing tools to aid integration of multiple datasets. It includes functions to create either a Seurat S4 object or SingleCellExperiment S4 object which are designed for single cell experiment analysis and contain slots for both expression data and metadata. The package provides function to import a raw count matrix file and a barcodes text file. The barcodes can be either be barcode file provided by Spatial Transcriptomics giving the coordinates of each probe or adjusted barcodes obtained by pre-processing the image using ST Spot Detector. The Scater R ^9^ package is used to calculate QC metrics for SingleCellExpeiment objects. Spaniel also provides a function to create a rasterised grob, compatible with ggplot2, from a pre-processed H&E image which is used as the background image for the plots. In this example, the sample is from the mouse olfactory bulb which is a multi-layered structure found in the forebrain of vertebrates (Figure1A).

**Figure 1:**
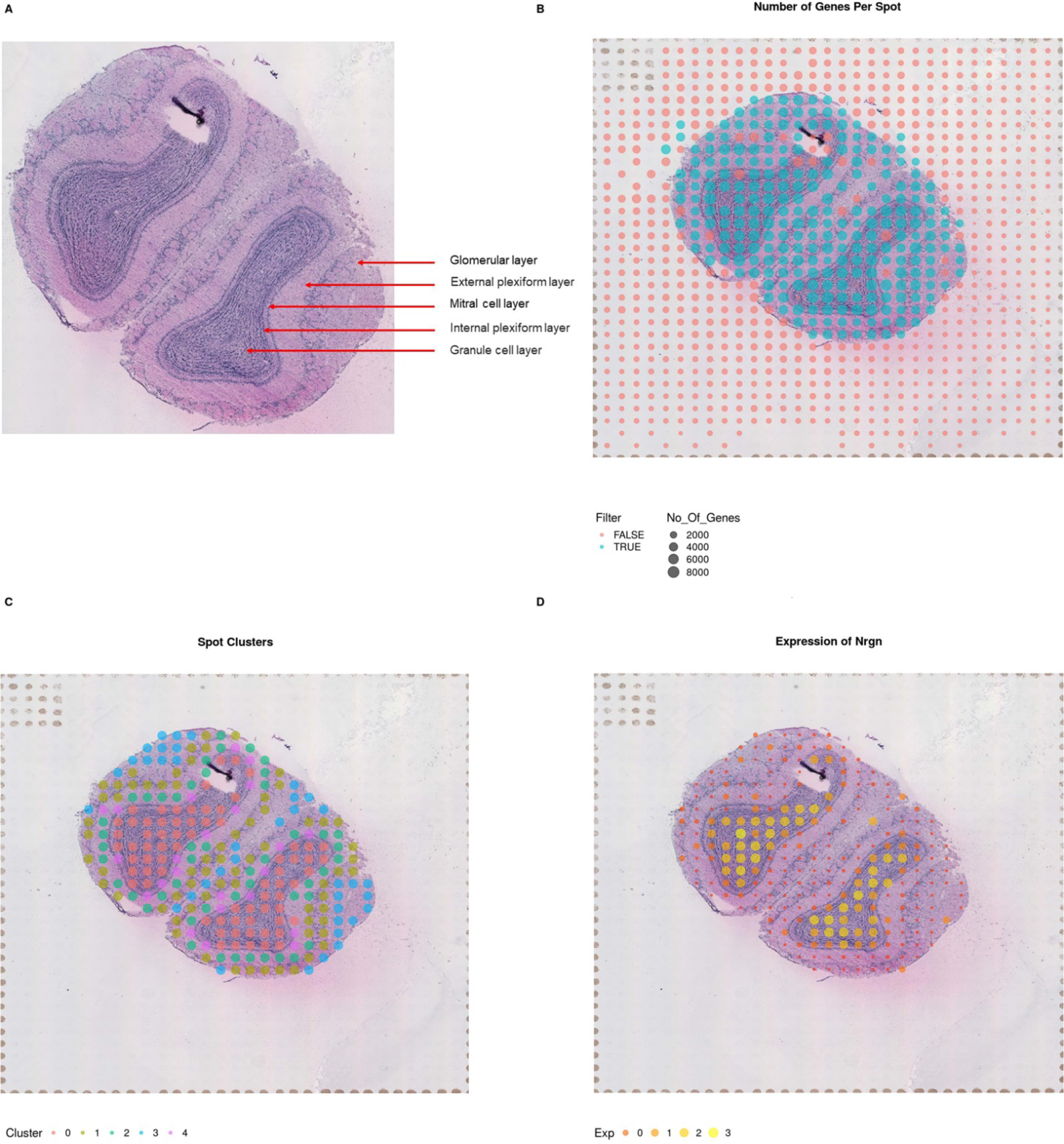
A) H&E image of mouse olfactory bulb showing the different layers. Example plots using Spaniel showing B) quality control metrics such as the number of genes detected per spot, C) the results of Seurat’s FindClusters clustering analysis, D) the expression of the *Nrgn* gene.

The ST_plot function provides a method to visualise metrics contained within the Seurat object overlaid onto the image of the tissue. This plot can be used initially as a quality control step as well as to visualise different clustering solutions or gene expression of specific genes.

Quality Control is a crucial first step in the analysis of this data. By inspecting the number of genes and number of reads per spot, using the ST_plot function, it is possible to pinpoint potential problems with the data that may confound downstream analysis. For example, spots which fall outside of the tissue with high number of counts may be an indicator of overall background RNA contamination. Figure 1B shows the number of genes per spot where spots with at least 2000 genes per spot are coloured in blue and spots with fewer than 2000 genes are coloured in red. The majority of spots above this threshold fall within the tissue area as expected, but a few fall outside the tissue area perhaps indicating contamination. One of the primary aims of Spatial Transcriptomics analysis is the identification of anatomical regions which share similar transcriptomes. As the package is tightly integrated with the Seurat package and SingleCellExperiment object it possible to choose a wide range of clustering methods such as the Seurat FindClusters function to identify regions sharing gene expression patterns. After clustering Seurat can be used to detect genes which are differentially expressed between clusters. The clusters, shown in figure 1C, appear to match the different layers of the olfactory bulb. For example, spots which fall into cluster 0 are detected in the granule cell layer. Through differential gene expression, *Nrgn* was found to be upregulated in cluster 0 compared to the other clusters (Figure 1D). This gene is neuron-specific and is the top gene associated with the granule cell layer of the olfactory bulb in the Allen Mouse Brain Atlas^10^.

Spaniel’s Shiny application can be hosted on a local computer or any service running R server (e.g. shinyserver.io) and offers the user 4 plots showing a) the number of genes detected per spot, b) the number of reads detected per spot, c) clustering results, d) the gene expression of a selected gene. The app uses a pre-processed S4 object and grob which are both saved in in RDS format allowing data to be transferred between the computational biologist performing the analysis and other researchers within the group. The gene plot allows users to select any individual gene to plot, whilst the cluster plot enables the visualisation of clustering results at multiple resolutions. Installation instruction and code can be found here: https://github.com/RachelQueen1/Spaniel

## References

1. Patrik L. Ståhl et al. Visualization and analysis of gene expression in tissue sections by spatial transcriptomics. Science 353, 78–82 (2016).

2. Zappia, L., Phipson, B. & Oshlack, A. Exploring the single-cell RNA-seq analysis landscape with the scRNA-tools database. PLoS Comput Biol 14, e1006245 (2018).

3. Fernandez Navarro, J., Lundeberg, J. & Stahl, P.L. ST viewer: a tool for analysis and visualization of spatial transcriptomics datasets. Bioinformatics 35, 1058–1060 (2019).

4. Butler, A., Hoffman, P., Smibert, P., Papalexi, E. & Satija, R. Integrating single-cell transcriptomic data across different conditions, technologies, and species. Nat Biotechnol 36, 411–420 (2018).

5. Lun A, R.D. SingleCellExperiment: S4 Classes for Single Cell Data. (2019).

6. Bolger, A.M., Lohse, M. & Usadel, B. Trimmomatic: a flexible trimmer for Illumina sequence data. Bioinformatics 30, 2114–2120 (2014).

7. Dobin, A. et al. STAR: ultrafast universal RNA-seq aligner. Bioinformatics 29, 15–21 (2013).

8. Anders, S., Pyl, P.T. & Huber, W. HTSeq--a Python framework to work with high-throughput sequencing data. Bioinformatics 31, 166–169 (2015).

9. McCarthy, D.J., Campbell, K.R., Lun, A.T. & Wills, Q.F. Scater: pre-processing, quality control, normalization and visualization of single-cell RNA-seq data in R. Bioinformatics 33, 1179–1186 (2017).

10. Lein, E.S. et al. Genome-wide atlas of gene expression in the adult mouse brain. Nature 445, 168–176 (2007).

